# Oculomotor Target Selection is Cortically Mediated by Complex Objects

**DOI:** 10.1101/2020.08.10.244319

**Authors:** Devin H. Kehoe, Jennifer Lewis, Mazyar Fallah

## Abstract

Successful oculomotor target selection often requires discriminating visual features but it remains contentious whether oculomotor substrates encoding saccade vectors functionally contribute to this process. One possibility is that visual features are discriminated cortically and oculomotor modules select the object with the highest activation in the set of all preprocessed cortical object representations, while an alternative possibility is that oculomotor modules actively discriminate potential targets based on visual features. If the latter view is correct, these modules should not require input from specialized visual cortices encoding the task relevant features. We therefore examined whether the latency of visual onset responses elicited by abrupt distractor onsets is consistent with input from specialized visual cortices by non-invasively measuring human saccade metrics (saccade curvature, endpoint deviations, saccade frequency, error proportion) as a function of distractor processing time for novel, visually complex distractors that had to be discriminated from a target to guide saccades. Visual onset response latencies were ~110 ms, consistent with projections from anterior cortical sites specialized for object processing. Surprisingly, oculomotor visual onset responses encoded features, as we manipulated the visual similarity between targets and distractors and observed an increased visual onset response magnitude and duration when the distractor was highly similar to the target, which was not attributable to an inhibitory processing delay. Visual onset responses were dynamically modulated by executive function, as these responses were anticipatorily extinguished over the time course of the experiment. As expected, the latency of distractor-related inhibition was modulated by the behavioral relevance of the distractor.

**Significance Statement:** We provide novel insights into the role of the oculomotor system in saccadic target selection by challenging the convention that neural substrates that encode oculomotor vectors functionally contribute to target discrimination. Our data show that the oculomotor system selects a winner from amongst the preprocessed object representations output from specialized visual cortices as supposed to discriminating visual features locally. We also challenge the convention that oculomotor visual onset responses are feature-invariant, as they encoded task-relevance.

## Introduction

When visual features define oculomotor targets (e.g., feature-based oddity search or template matching), successful target selection requires discriminating on the basis of features. The visual features characterizing potential oculomotor targets modulate activity in the intermediate layers of the superior colliculus (SCi) (Horwitz and Newsome, 2001; McPeek and Keller, 2002; Shen and Paré, 2012) and frontal eye fields (FEF) (Bichot and Schall, 1999; Thompson et al., 2005) visuomotor (VM) cells. Inactivation of SCi causes feature-based target selection deficits for saccades (McPeek and Keller, 2004) and manual button presses (Lovejoy and Krauzlis, 2010). Conversely, subthreshold microstimulation of SCi or FEF biases feature-based target selection for eye movements (Carello and Krauzlis, 2004; Dorris et al., 2007; McPeek et al., 2003; McPeek, 2006) and facilitates perceptual discriminations (Cavanaugh and Wurtz, 2004; Müller et al., 2005; Moore and Fallah, 2001, 2004) by modulating downstream cortical visual representations (Moore and Armstrong, 2003; Monosov et al., 2011). These observations demonstrate that SCi and FEF are necessary for feature-based target selection, which guides both action and perception, but it remains unclear whether SCi and FEF contribute to feature-based target selection by selecting the feature-weighted cortical object representation with the highest activation or by actively resolving competition between cortical object representations by locally discriminating features.

Given that neither the SCi nor FEF possess any inherent feature selectivities but encode the feature-based behavioral relevance of potential saccade targets (Boehnke and Munoz, 2008; Fecteau and Munoz, 2006) and are richly interconnected with the visual cortices (Schall et al., 1995; White and Munoz, 2011), it seems likely that SCi and FEF select a target based on the available feature-based cortical visual representations. This view suggests that SCi and FEF are necessary but insufficient for feature-based target selection, as they lack the requisite specificity for discrimination. If SCi or FEF were sufficient for feature-based target selection, they should not require input from visual cortices specialized for processing task-relevant visual features. Here, we examine whether the timing of visual onset responses in saccade vector-encoding oculomotor substrates is consistent with feedforward projections from cortical modules specialized for processing relevant stimulus features.

Kehoe and Fallah (2017) developed a non-invasive method for estimating the latency of oculomotor excitation and suppression during target selection by measuring distractor-elicited saccade curvature as a function of time between an abrupt, task-irrelevant distractor onset and saccade execution. Targets onset prior to distractors allowing us to continuously analyze the entire time course of distractor processing. The theoretical foundation of this non-invasive technique is rooted in the spatiotemporal oculomotor interactions in SCi and FEF associated with curved saccades: during target selection, when there is transient excitation (McPeek et al., 2003; McPeek, 2006; Port and Wurtz, 2003) or suppression (White et al., 2012) of VM cells encoding a distractor ~20-30 ms prior to saccade execution, saccades curve towards or away (respectively) from the distractor and the magnitude of the excitation or suppression in this epoch is proportional to the magnitude of saccade curvatures. In the excitatory case, this mechanism has been causally demonstrated with microstimulation (McPeek et al., 2003; McPeek, 2006).

Kehoe and Fallah (2017) measured an ~20 ms difference between the latencies of peak excitation elicited by luminance- and color-modulated task-irrelevant distractors, consistent with the transient visual onset burst latency differences between similar stimuli observed in collicular VM cells (White et al., 2009) and between cells in the magno-/parvocellular or dorsal/ventral processing streams in early visual modules (Nowak and Bullier, 1997; Schmolesky et al., 1998). We also observed a peak excitation latency of ~115 ms and reasoned that since transient distractor activity must occur ~30 ms prior to saccade execution to elicit saccade curvature (McPeek et al., 2003; McPeek, 2006; Port and Wurtz, 2003; White et al., 2012) and since transient visual onset bursts in collicular VM cells reach peak levels of activity ~30 ms after their onset (McPeek and Keller, 2002), oculomotor onset bursts occurred ~55 ms after distractor onset on average, as 115 ms – 30 ms – 30 ms = 55 ms, consistent with collicular visual onset burst latencies (reviewed by Boehnke and Munoz, 2008). Such short latencies are consistent with distractor representations projected into oculomotor substrates by early cortical visual modules, as expected given the low-level visual features of the distractors.

In the current study, we modified our technique such that subjects performed a template-matching visual discrimination between complex, novel target and distractor objects and executed saccades to the target, as these stimuli would likely require cortical feature processing in late stages of the ventral visual processing stream. As in Kehoe and Fallah (2017), we randomized the interval between distractor and target onsets (distractor-target onset asynchrony [DTOA]) but ensured feature processing of the stimuli by presenting the distractor before the target on 50% of trials so that the order of stimulus onset did not provide target information. As such, we analyzed saccade curvature as a function of time between saccade initiation and distractor onset for correct trials in which the distractor onset before the target. Additionally, we examined saccade endpoint deviations, saccade frequencies, and error proportions, as they are also indicative of oculomotor excitatory and suppressive processing: (a) subthreshold microstimulation of a secondary saccade vector in SCi or FEF concurrent with saccade initiation causes saccade endpoints to shift towards the stimulated vector and the magnitude of these deviations is proportional to the intensity of the microstimulation (Fuchs and Robinson, 1969; Glimcher and Sparks, 1993; Robinson, 1972). Conversely, saccade endpoints deviate away from the site of inhibitory collicular injections (Lee et al., 1988). (b) Collicular stimulation transiently elicits rapid (~5 ms) inhibition in surrounding loci (Munoz and Istvan, 1998) lowering the behavioral likelihood of saccade initiation (Reingold and Stampe, 2002). (c) Target selection is guided by the available representations in SCi, as an inhibitory injection at the target locus greatly impairs accurate target selection (McPeek and Keller, 2004). We also manipulated the visual similarity between targets and distractors to examine when behavioral relevance encoding of stimuli emerged during the oculomotor processing time course. Based on previous investigations of feature-based saccadic target selection, we expected that the initial excitatory response would be feature invariant, while the subsequent suppressive response would vary in latency and magnitude between similarity conditions (Fecteau and Munoz, 2006). Finally, to examine whether perceptual learning modulates the visual onset response, we examined whether distractor processing was similar across the time course of the experimental session.

## Methods

### Participants

36 York University undergraduate students (17-30 years old, 6 male) participated in the experiment for course credit. Participants had normal or corrected-to-normal visual acuity and were naïve to the purpose and design of the experiment. Informed consent was obtained prior to participation. All research was approved by York University’s Human Participants Review Committee.

### Stimuli

6 stimuli used in a previous experiment (Kehoe et al., 2018) were constructed offline using MATLAB (MathWorks, Natick, MA) by intersecting 6 or 7 vertical and horizontal line segments (1°× 0.08°) together at right angles in a configuration that did not resemble meaningful alphanumerical characters to an English speaker (similar to Palmer, 1978; see Figure 1*A*). Individual line segments occupied 1 of 12 possible locations that were embedded in an imaginary box (2° × 2°). All stimuli were linearly related to one another in the number of line segments differences between them. Stimuli were assigned to an integer-valued location on a conceptual number line in which the absolute difference in number line position between any two stimuli was equal to the number of line segment differences between them and is therefore inversely proportional to their visual similarity. The difference in number line position between stimuli is herein referred to as objective similarity (OS). The 6 stimuli were divided into 3 subsets of 2 stimuli: {3,4}, {2,5}, and {1,6}. Each subset was characterized by a unique OS value: OS = 1, OS = 3, and OS = 5, referred to herein as OS1, OS3, and OS5 (respectively). The stimuli were white (CIE*xy* = [.29, .30], luminance = 126.02 cd/m^2^) and were displayed against a black (CIE*xy* = [.27, .26], luminance = 0.20 cd/m^2^) background on a 21-inch CRT monitor (85 Hz, 1024 × 768). Participants viewed stimuli in a dimly lit room from a viewing distance of 57 cm with a headrest stabilizing their head position.

**Figure 1.**
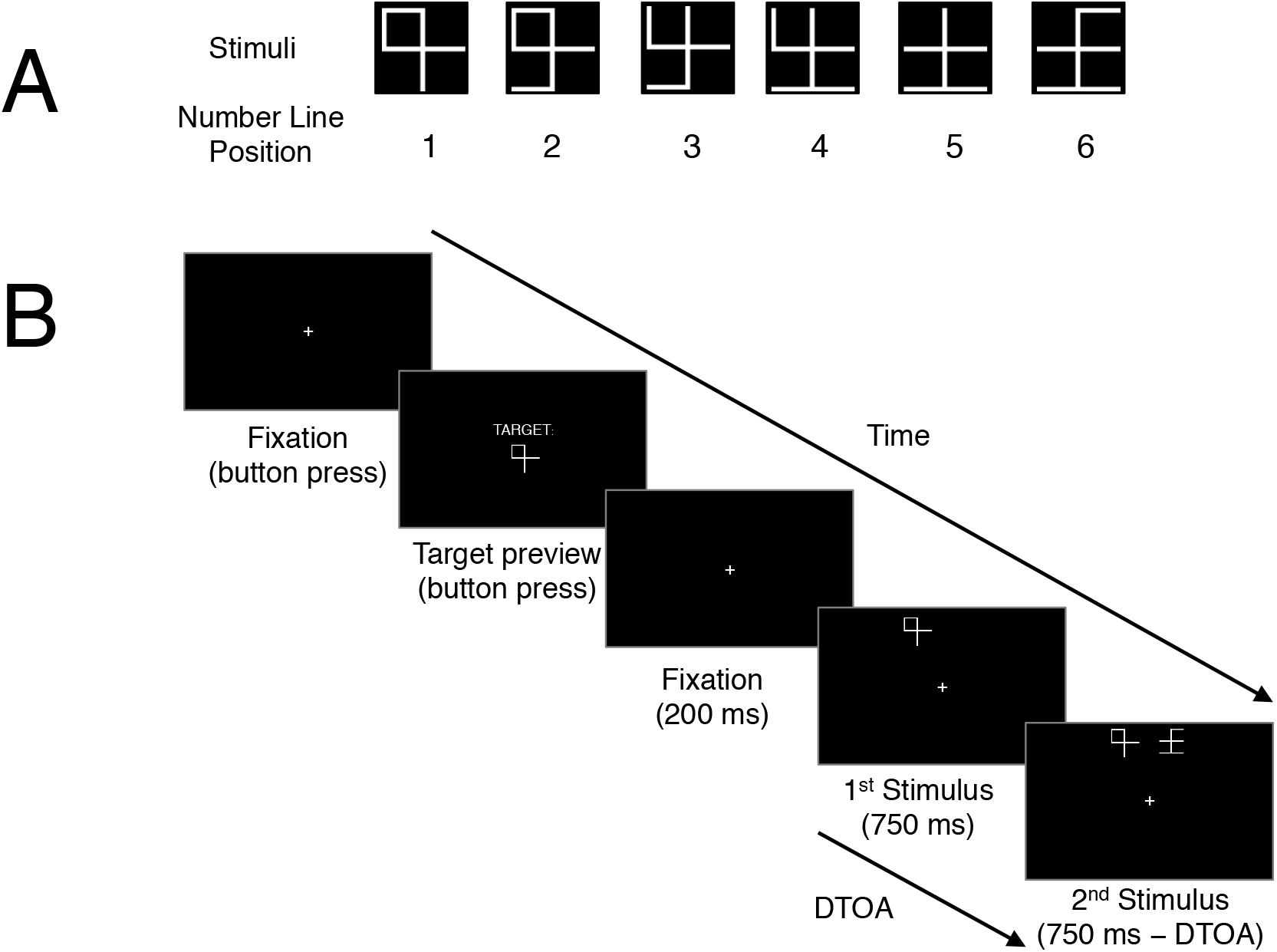
Discrimination saccade task stimuli and displays. *A*: Stimuli used in the current experiment. Stimuli were placed on an integer-valued conceptual number line in which the absolute difference between number line positions corresponds to the number of line segment differences, referred to as objective similarity (OS). *B*: Trial structure. Participants pressed a button to preview the target stimulus until they were familiar with it. Participants then pressed the button again to initiate the discrimination display. After maintaining fixation for 200 ms, the display was presented for 750 ms or until a saccade to one of the stimuli was detected. The target and distractor onsets were separated by a randomized interval, referred to as distractor-target onset asynchrony (DTOA).

### Apparatus and Measurement

Stimulus presentation was controlled using a computer running Presentation software (Neurobehavioral Systems, Berkeley, CA). Manual button responses were collected on a serial response box (RB-540; Cedrus, San Pedro, CA). Eye position was recorded using infrared eye tracking (500 Hz, EyeLink II; SR Research, Ontario, Canada). The eye tracker was calibrated using a nine-point grid at the beginning and halfway point of each experimental session, and as needed. All data processing and statistical analysis was performed using MATLAB.

### Task Procedure

Trials were initiated by button pressing (see Figure 1*B*). A target preview was presented until participants pressed the button a second time. A white fixation cross (0.4°× 0.4°) then appeared in the center of the display. After participants maintained fixation (1.89° × 1.89° window) for 200 ms, the target and distractor appeared at 1 of 4 locations. The target and distractors were always in the same vertical hemifield, were equidistant from fixation (8° eccentricity), and angularly separated from the vertical meridian by 22.5°. The relative time between distractor and target onsets (distractor-target onset asynchrony [DTOA]) was −150, −100, −50, 0, 50, 100, or 150 ms in which a positive value indicates that the target onset first. To correctly discriminate the target, participants were instructed to maintain fixation, use their peripheral vision to determine which stimulus was the target, and then make a saccade to it. The trial ended when a saccade was made to the target (correct) or distractor (incorrect) or 750 ms had elapsed from the time of the first stimulus onset (time-out). An error tone and message were used to indicate incorrect and time-out trials. Time-out trials were randomly replaced back into the block. Participants received accuracy feedback at the end of each block.

Participants completed 1 session with 6 blocks of 75 trials for a total of 450 trials. Each block contained 3 repeats of 25 conditions (3 OS conditions × 7 DTOA conditions + 4 no distractor baseline trials, one for each stimulus location) in randomized order. On each trial, the hemifield (upper vs. lower), left/right order of the target and distractor, and target/distractor stimulus assignment (e.g., target = 1, distractor = 6 vs. target = 6, distractor = 1) was randomized with equal probability.

### Saccade Detection

Saccades were detected, visualized, filtered and analyzed offline using customized MATLAB algorithms. Saccades were defined as a velocity exceeding 20 °/s for at least 8 ms and a peak velocity exceeding 50 °/s. Saccadic reaction time (SRT) was defined as the time from target onset to saccade initiation. Trials that contained blinks (1.98%), corrective saccades (1.01%), saccade amplitudes < 1° (1.75%), endpoint deviations > 3° from the center of the target (5.26%), fixation drifts > 0.5° during the pre-saccadic latency period (4.27%), or an SRT < 100 ms (3.49%) were excluded from further analysis leaving 82.24% of the data remaining.

Saccade curvatures were quantified as the sum of all orthogonal deviations from a straight line between the start and end of the saccade in degrees visual angle. Endpoint deviations were quantified as the angular separation between the saccade endpoint and the center of the target in polar degrees. Mean baseline saccade curvature and endpoint deviation for each participant at each target location was subtracted from the data to reduce idiosyncratic movement. These metrics were coded so that positive values correspond to deviations towards the distractor, while negative values correspond to deviations away from the distractor.

### Data Analysis *Task Performance*

To ensure that the task was performed at above chance levels, a binomial exact test was conducted on the frequency of correct trials for each participant. Participants who did not score above chance (*p* < .05) were removed from subsequent analyses.

### Distractor Processing Time

We only analyzed trials on which the target onset prior to or synchronously with distractor onset (DTOA ≥ 0). Time after distractor onset was computed as the time between distractor onset and saccade initiation. We fit a Gaussian kernel smoother with a bandwidth of 10 ms to the saccade curvature and endpoint deviation data from each subject as a function of time after distractor onset. A sliding two-tailed, Wilcoxon signed-rank test was used to determine the distractor processing epochs at which saccade curvature and endpoint deviation were significantly different than zero. Similarly, a Gaussian kernel density estimator (KDE) with a bandwidth of 10 ms was fit to the distributions of correct and erroneous saccades for each subject to estimate saccade frequency and error proportion as a function of time after distractor onset. Saccade frequencies appeared to be distributed as an exponentially-modified Gaussian (EMG) function of time after distractor onset but for a transient decrease in the range of ~125-250 ms. To estimate the distractor processing time (*t*) of this transient decrease, we first fit an expectation EMG model to the data outside of this range. The model was defined as

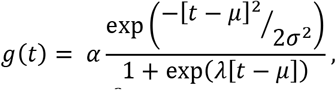

where *α* scales the height of the model, *μ* and *σ*^2^ are the mean and variance parameters for the Gaussian component of the model, and *λ* scales the skew of the model. We fit the model by maximizing the log-likelihood of the parameters using the normal distribution. We validated the model with a ratio test comparing the fit to a one-parameter null model. Second, a sliding one-tailed Wilcoxon signed-rank test was used to determine the distractor processing time that saccade frequencies were significantly lower than the expectation model. A sliding Wilcoxon signed-rank test was used to determine the distractor processing epochs at which error proportion was significantly greater than the standard error of the proportion. We only analyzed saccade curvature, endpoint deviation, and error proportion for distractor processing times with at least 20 saccades, i.e., 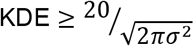, where *σ* = 10 (KDE bandwidth).

This analysis was repeated to examine (a) the overall distractor processing time course; (b) distractor processing time course differences between OS conditions; and (c) distractor processing time course differences between the first, second, and final 1/3 of experimental blocks. All saccade metrics were compared between conditions with a sliding two-tailed, Wilcoxon signed-rank test to determine when processing was significantly different between conditions. For all sliding inferential analyses, epochs were considered significant when *p* < .05 for at least 10 ms. Significant epochs separated by ≤ 5 ms were pooled together. Bonferroni multiplicity corrections were used for all multiple inferential comparisons.

### Disentangling SRT and Distractor Processing Time

We performed several analyses to ensure that any potential effects of distractor processing time are not confounded by systematic SRT differences between conditions. First, we examined potential SRT mean differences between DTOA (0, 50, 100, 150), OS (OS1, OS3, OS5), and experimental block (early, middle, late) conditions using a linear mixed-effects model with fixed effects for all conditional main effects and interactions and with random subject intercepts for each fixed effect similar to repeated-measures ANOVA. A marginal *F*-test with planned orthogonal comparisons as *post hocs* was used to analyze all fixed effects. To examine potential speed-accuracy trade-offs, we repeated this analysis for error proportions with a generalized linear mixed-effects model with a logit link function to the binomial distribution. Next, we repeated our data smoothing analysis of saccade metrics as a function of distractor processing time (see section “Distractor Processing Time” above) separately across DTOA conditions. This allowed us to investigate whether excitatory activity is related to distractor processing time or SRT, as SRTs vary across DTOA for a fixed distractor processing time. For example, 100 ms of distractor processing time could arise from a synchronous target-distractor onset (DTOA0) and an SRT of 100 ms or from a distractor onset 150 ms after the target (DTOA150) and an SRT of 250 ms. To analyze the distractor processing epochs with differences between DTOA conditions, we used a sliding Friedman test with the same inferential conventions as in the previous section.

## Results

All participants correctly discriminated the target above chance (all *ps ≤* .009), except for one participant (*p* = .050) who was therefore removed from subsequent analyses. The group mean percentage of correct target discriminations was 89.67% *(SE* = 0.92%).

### Overall Distractor Processing

We analyzed the behavioral effects of distractor processing time averaged across all conditions (see Figure 2). For saccade curvature, there was an initial epoch of gradually decreasing excitatory processing (0-69 ms), followed by a transient excitatory epoch (133-168 ms), and then an extended epoch of inhibitory processing (247-500 ms) (see Figure 2*A*). Endpoint deviation showed an extended excitatory epoch (108-243 ms) (see Figure 2*B*). The expectation model provided a good fit to saccade frequency (*p* < .001) and indicated that there was a transient drop in saccade frequency (130-230 ms) (see Figure 2*C*). There was an abrupt onset of erroneous saccades that gradually decreased over distractor processing time (154-349 ms) (see Figure 2*D*).

**Figure 2.**
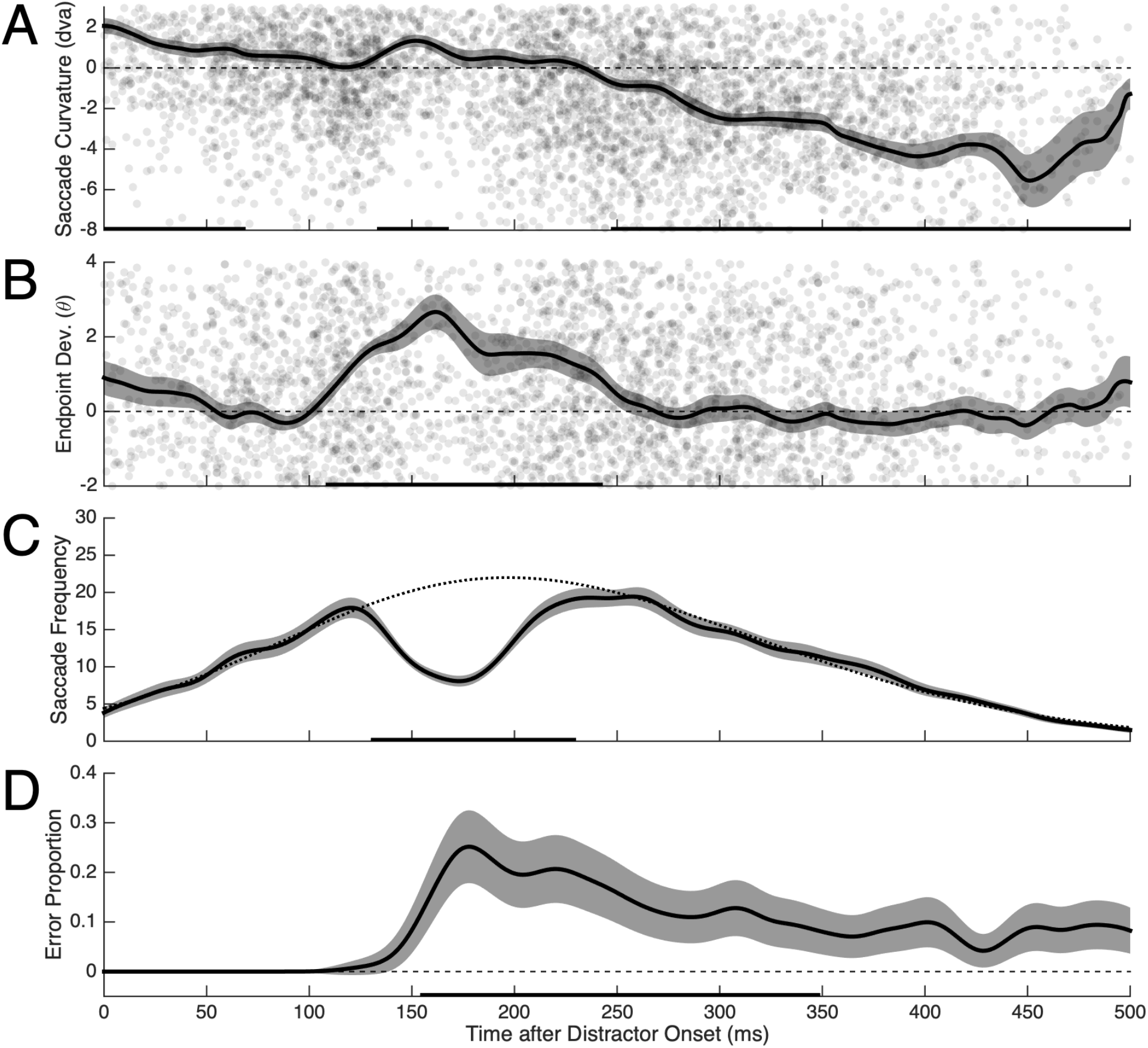
Smoothed overall saccade metrics as a function of distractor processing time. Shaded error bars represent standard error. All statistical analyses were considered significant at *p* < .05. *A, B*: Data points indicate individual saccades. Black bars along the abscissa indicate significant differences from zero (two-tailed Wilcoxon signed-rank test). *A*: Mean saccade curvature. *B*: Mean endpoint deviation. *C*: Sum saccade frequency. Dashed line indicates fitted exponentially-modified Gaussian expectation model. Black bars along the abscissa indicate significant differences below expectation (one-tailed Wilcoxon signed-rank test). *D*: Proportion of errors. Black bars along the abscissa indicate significant differences above the standard error of the proportion (one-tailed Wilcoxon signed-rank test).

### OS Condition Distractor Processing Differences

We analyzed the behavioral effects of distractor processing time separately across OS conditions (see Figure 3). For saccade curvature, there was initial, excitatory processing inconsistently observed across conditions. Critically, there was a transient excitatory epoch observed across all OS conditions and the duration of this epoch was related to OS (OS1: 144-156 ms; OS3: 132-167 ms; OS5: 134-195 ms). Similarly, the response magnitude of excitatory activity in this epoch was related to OS (OS5>OS1: 152-198 ms). There was also a secondary transient excitatory epoch uniquely observed in the OS1 condition (218-234 ms) in which OS1>OS3 (213-235 ms) and OS1>OS5 (203-230 ms). Finally, there was an extended inhibitory epoch that onset first in the OS3 (245-461 ms) and OS5 (247-478 ms) conditions, and later in the OS1 condition (303-482 ms). OS3 and OS5 saccade curvature was more negatively curved than OS1 curvature during the inhibitory delay in the OS1 condition (OS1>OS3: 271-310 ms; OS1>OS5: 269-336 ms). There was a brief period of negative curvature in the OS3 condition preceding the extended inhibitory epoch (215-229 ms), which we think reflects the true inhibitory onset in the OS3 condition given both the trend of the data and the observation that the sliding inferential analysis was marginally significant (.051 ≤ *p ≤* .109) throughout the interval separating the brief and extended inhibitory epochs (216-244 ms).

**Figure 3.**
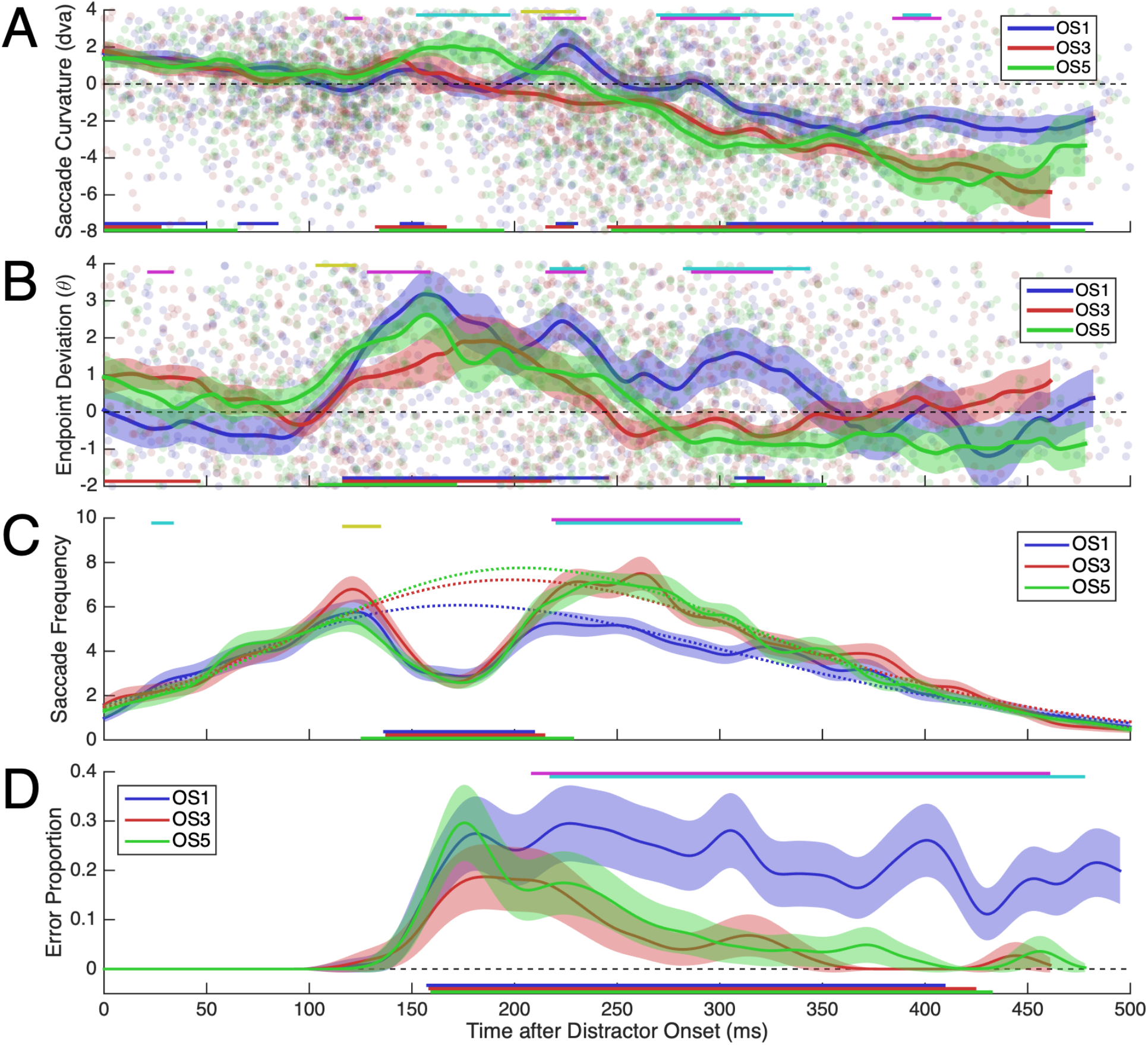
Smoothed saccade metrics as a function of distractor processing time in the OS1 (blue), OS3 (red), and OS5 (green) conditions. Shaded error bars represent standard error. Colored bars above panels indicate significant differences between conditions (OS1 vs. OS3 = magenta; OS1 vs. OS5 = cyan; OS3 vs. OS5 = yellow; two-tailed Wilcoxon signed-rank test). All statistical analyses were considered significant at *p* < .05. *A*, *B*: Data points indicate individual saccades. Colored bars along the abscissa indicate significant differences from zero (two-tailed Wilcoxon signed-rank test). *A*: Mean saccade curvature. *B*: Mean endpoint deviation. *C*: Sum saccade frequency. Dashed lines indicate fitted exponentially-modified Gaussian expectation models. Colored bars along the abscissa indicate significant differences below expectation (one-tailed Wilcoxon signed-rank test). *D*: Proportion of errors. Colored bars along the abscissa indicate significant differences above the standard error of the proportion (one-tailed Wilcoxon signed-rank test).

Endpoint deviation also indicated an excitatory epoch which onset first in the OS5 condition (104-172 ms) and shortly thereafter in the OS1 (116-246 ms) and OS3 conditions (116-218 ms) (see Figure 3*B*). As such, OS5 endpoint deviation was higher than OS3 deviation during this delay (OS5>OS3: 103-123 ms). Critically, during the excitatory epoch, endpoint deviation was also higher in the OS1 condition than in OS5 (OS1>OS3: 128-159 ms). OS1 endpoint deviation was higher than in OS3 and OS5 conditions concurrent with the secondary excitatory epoch observed for saccade curvature in the OS1 condition (OS1>OS3: 215-235 ms; OS1>OS5: 217-234 ms). There was also a secondary excitatory epoch in the OS1 (196-244 ms) and OS5 conditions (196-224 ms). Endpoint deviations were otherwise non-significant but for an additional excitatory epoch in the OS1 condition in which OS1 endpoint deviation was greater than OS3 and OS5 endpoint deviation (OS1>OS3: 286-326 ms; OS1>OS5: 282-344 ms). Interestingly, this epoch somewhat corresponded to the inhibitory delay period between the OS1 condition vs. OS3 and OS5 conditions observed with saccade curvature.

The expectation model provided a good fit to saccade frequency across OS conditions (all *p* < .001) (see Figure 3*C*). There was a transient drop in the likelihood of making a saccade across OS conditions (OS1: 136-210 ms; OS3: 137-215 ms; OS5: 125-229 ms). Interestingly, there was a lower frequency of saccades in the OS1 condition than both the OS3 and OS5 conditions (OS1>OS3: 218-310 ms; OS1>OS5: 220-311 ms), coinciding with the secondary excitatory epoch observed uniquely in the OS1 condition and also coinciding with the inhibitory delay observed in the OS1 condition relative to the OS3 and OS5 conditions.

There was a transient increase in error rates consistent across OS conditions (OS1: 157-410; OS3: 158-425 ms; OS5: 159-433 ms) (see Figure 3*D*). Errors were consistently higher in the OS1 condition than the OS3 and OS5 conditions (OS1>OS3: 208-461 ms; OS1>OS5: 217-478 ms).

### Experimental Block Distractor Processing Differences

Examining the effects of distractor processing time separately over the early (E), middle (M), and late (L) experimental blocks of the experiment demonstrated five notable effects (see Figure 4): (a) The saccade curvature excitatory response was longest in the early blocks (124-168 ms), reduced in the middle blocks (142-163 ms), and was completely extinguished in the late blocks (see Figure 4*A*). (b) The latency of the saccade curvature inhibitory response decreased over the course of the experiment (E: 276 ms; M: 251 ms; L: 240 ms). (c) Endpoint deviation in the initial excitatory epoch was considerably higher in the early blocks than later in the experiment (E>M: 161-198 ms; E>L: 166-189 ms) (see Figure 4*B*). (d) The latency of the transient drop in saccade likelihood was consistent across experimental blocks (E: 128 ms; M: 130 ms; L: 133 ms; all expectation models *p* < .001) (see Figure 4*C*). (e) Errors during the initial excitatory epoch were higher in the early blocks than the late (E>L: 123-154 ms) blocks (see Figure 4*D*).

**Figure 4.**
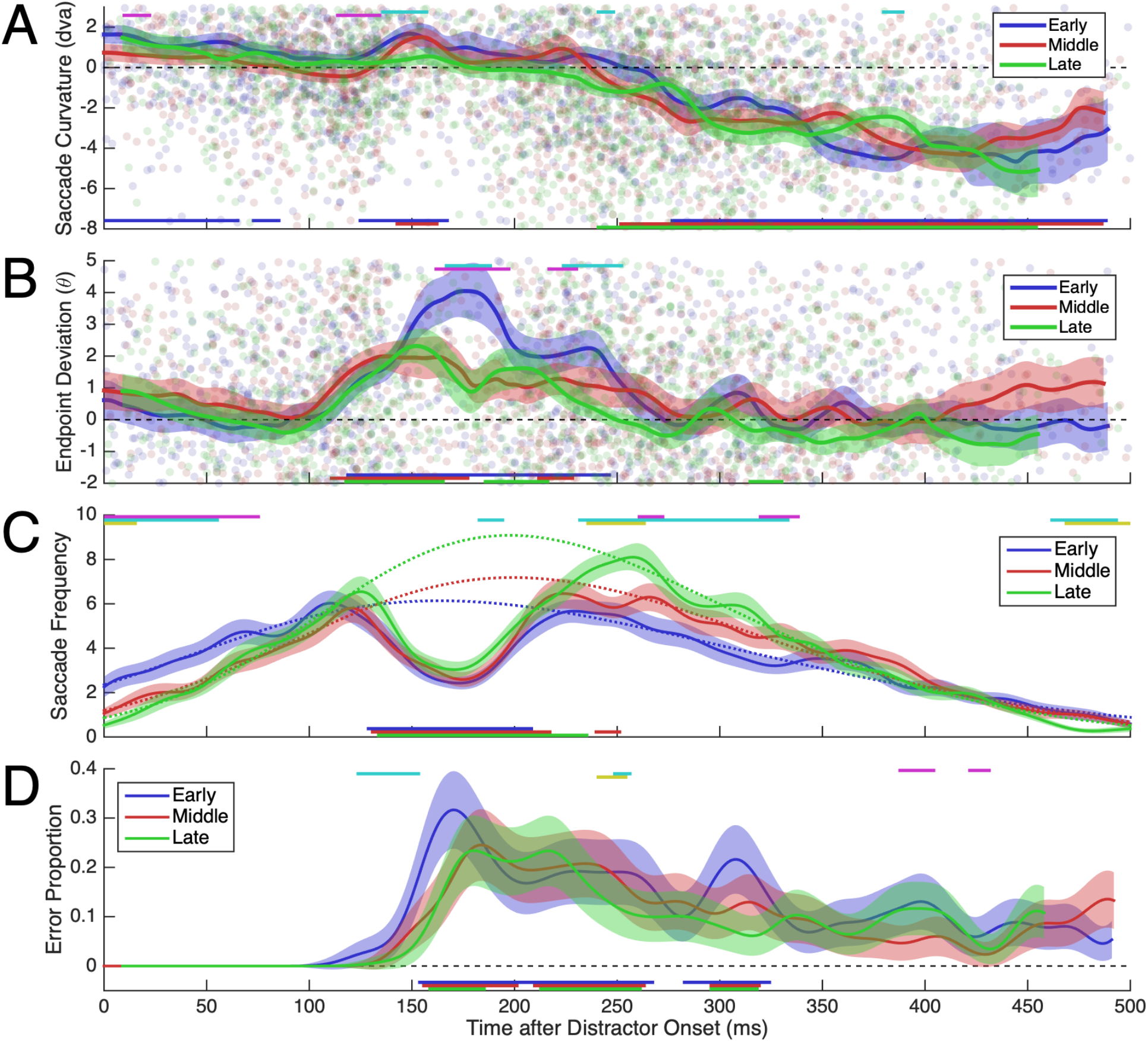
Smoothed saccade metrics as a function of distractor processing time in the early (blue), middle (red), and late (green) experimental blocks. Shaded error bars represent standard error. Colored bars above panels indicate significant differences between conditions (early vs. middle = magenta; early vs. late = cyan; middle vs. late = yellow; two-tailed Wilcoxon signed-rank test). All statistical analyses were considered significant at *p* < .05. *A,B*: Data points indicate individual saccades. Colored bars along the abscissa indicate significant differences from zero (two-tailed Wilcoxon signed-rank test). *A*: Mean saccade curvature. *B*: Mean endpoint deviation. *C*: Sum saccade frequency. Dashed lines indicate fitted exponentially-modified Gaussian expectation models. Colored bars along the abscissa indicate significant differences below expectation (one-tailed Wilcoxon signed-rank test). *D*: Proportion of errors. Colored bars along the abscissa indicate significant differences above the standard error of the proportion (one-tailed Wilcoxon signed-rank test).

### SRT Processing Differences

There were no SRT main effects or interactions between conditions (all *F* ≤ 1.50, all *p* ≥ .127) (see Figure 5). There was a significant effect of DTOA on error proportions (*F*[3,1218] = 37.37, *p* < .001). All pairwise differences were significant (all *F* ≥ 4.76, all *p* ≤ .029), except for DTOA150 and DTOA100 (*F*[1,1218] = 2.59, *p* = .108). There was a significant effect of OS on error proportions (*F*[2,1218] = 51.28, *p* < .001). All pairwise differences were significant (all *F* ≥ 55.73, all *p* < .001), except for OS3 and OS5 (*F*[1,1218] = 3.40, *p* = .065). There were no other error proportion main effects or interactions between conditions.

**Figure 5.**
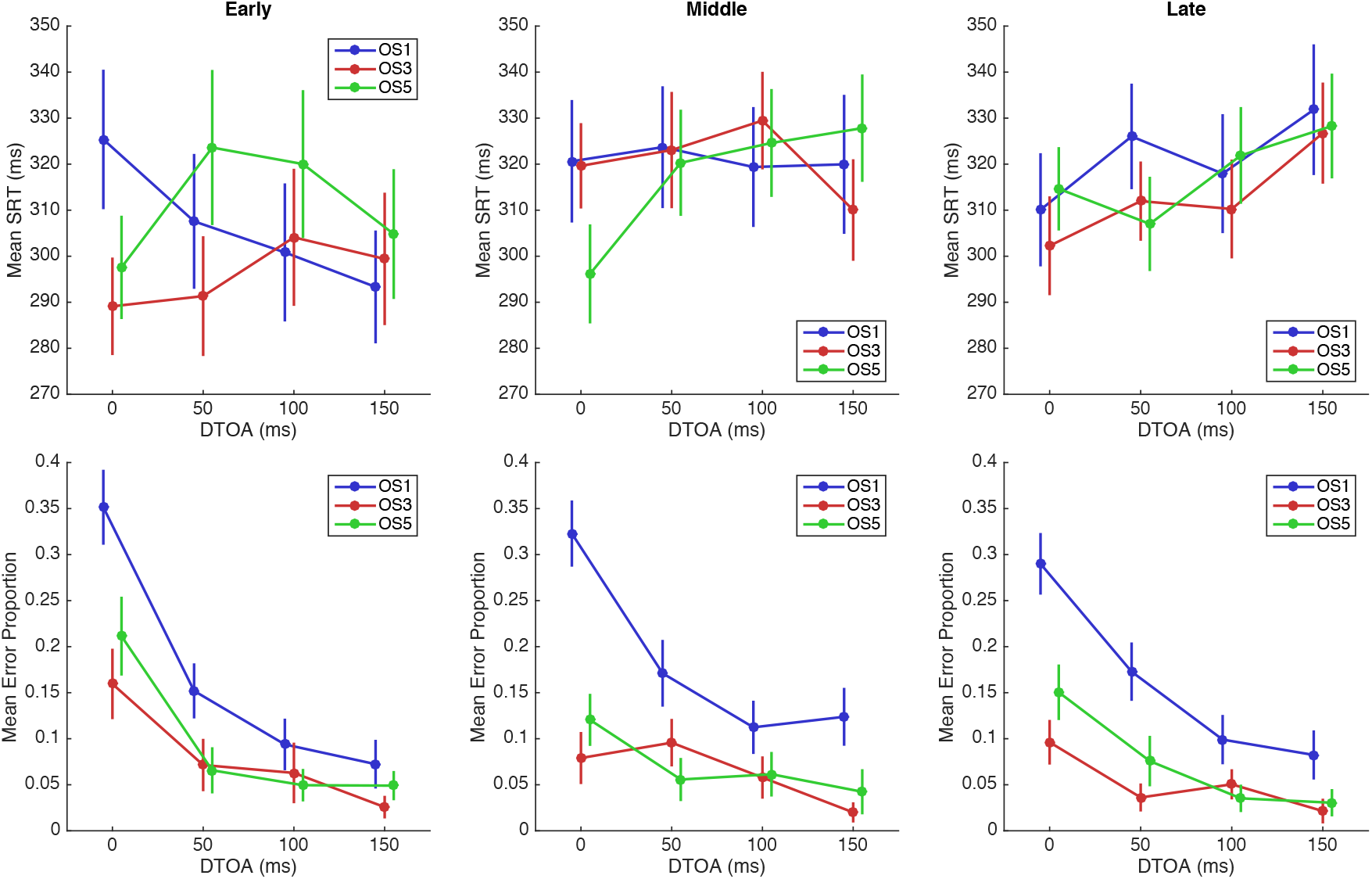
Mean SRT (top row) and error proportion (bottom row) differences between DTOA, OS, and experimental block conditions. DTOA is plotted along abscissa. OS conditions are color-coded (OS1 = blue, OS3 = red, OS5 = green). Experimental block conditions are plotted by column (early = left, middle = middle, late = right). Error bars represent standard error.

We analyzed the behavioral effects of distractor processing separately across DTOA conditions (see Figure 6). There were effects of both SRT and distractor processing time on saccade curvature (see Figure 6*A*). Across DTOA conditions, there were consistent positive curvatures after ~140 ms of distractor processing (DTOA0: 139-229 ms; DTOA50: 145-181 ms; DTOA100: 138-159 ms), except in the DTOA150 condition in which the excitatory response failed to meet the 10 ms criterion but was significantly greater than zero for 9 ms (140-148 ms). During the excitatory epoch, there were differences between DTOA conditions (139-220 ms), likely driven by higher saccade curvature in the DTOA0 condition than the remaining conditions. These results indicate that the magnitude of excitatory responses were modulated my DTOA condition, but critically, not the timing. There was also positive curvature clearly associated with SRTs, as positive curvature was observed in ~50 ms intervals across DTOA conditions, which corresponded to SRTs of ~120-140 ms in each condition (DTOA0: 139-229 ms + 0 ms = SRT 139-229 ms; DTOA100: 18-82 ms + 100 ms = SRT 118-182 ms; DTOA150: −24-33 ms + 150 ms = SRT 126-183 ms). In fact, in the DTOA150 condition, this corresponded to positive curvature prior to the distractor onset. This SRT-related positive curvature was not significant in the DTOA50 condition, but still qualitatively exhibited the same trend. The onset of inhibition did vary somewhat across DTOA conditions as a function of distractor processing time (DTOA0: 257 ms; DTOA50: 265 ms; DTOA100: 233 ms; DTOA150: 277 ms), but there were no saccade curvature differences between DTOA conditions in the inhibitory epoch.

**Figure 6.**
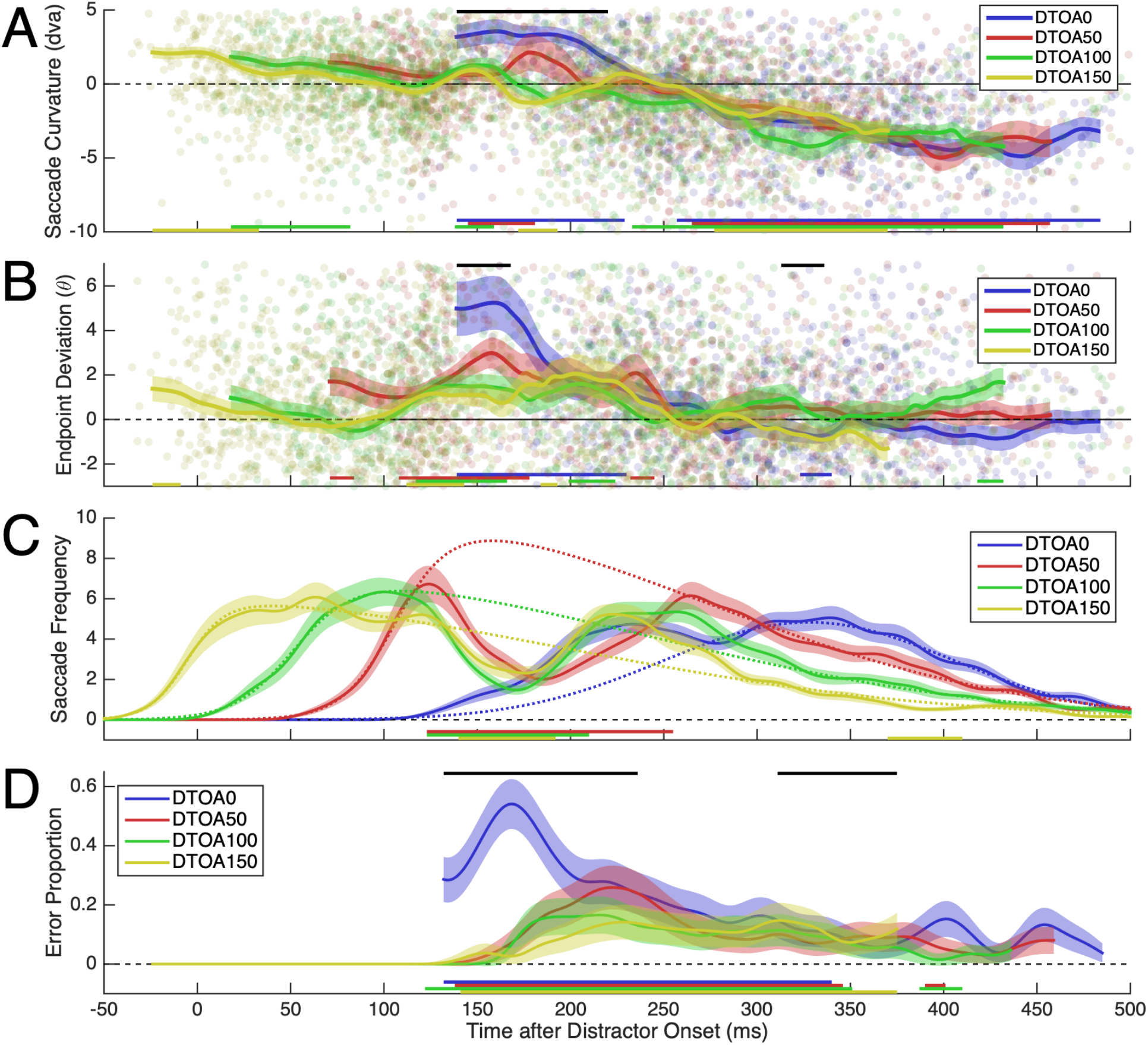
Smoothed saccade metrics as a function of distractor processing time in the DTOA0 (blue), DTOA50 (red), DTOA100 (green), and DTOA150 (yellow). Shaded error bars represent standard error. Black bars above panels indicate significant differences between conditions (Friedman test). All statistical analyses were considered significant at *p* < .05. *A, B:* Data points indicate individual saccades. Colored bars along the abscissa indicate significant differences from zero (two-tailed Wilcoxon signed-rank test). *A*: Mean saccade curvature. *B*: Mean endpoint deviation. *C:* Sum saccade frequency. Dashed lines indicate fitted exponentially-modified Gaussian expectation models. Colored bars along the abscissa indicate significant differences below expectation (one-tailed Wilcoxon signed-rank test). *D:* Proportion of errors. Colored bars along the abscissa indicate significant differences above the standard error of the proportion (one-tailed Wilcoxon signed-rank test).

Endpoint deviation was weakly related to SRT (DTOA50: 71-84 ms; DTOA150: −24-(−9) ms) (see Figure 6*B*). Visual onset response latencies were related to distractor processing time and were consistent with the overall estimates (DTOA50: 108 ms; DTOA100: 117 ms; DTOA150: 112 ms). However, there were insufficient observations in the DTOA0 condition to determine if the visual onset response latency was as early as ~110 ms of distractor processing. As with saccade curvature, there were differences between DTOA conditions during the excitatory epoch (139-168 ms), suggesting that the magnitude of excitatory responses were modulated my DTOA condition, but critically, not the timing.

Saccade frequency distributions were offset by ~50 ms between DTOA conditions, as expected since SRT was measured from target onset (see Figure 6*C*). However, across DTOA conditions, there was a drop in saccadic likelihood with latencies related to distractor processing time (DOTA50: 123 ms; DTOA100: 123 ms; DTOA150: 140 ms), except for DTOA0 in which there were too few observations in this range (all expectation models *p* < .001). As expected, the expectation model failed to reproduce the DTOA0 saccade frequency distribution in the 125-250 ms range, as it was fit outside of this range.

Error onsets varied somewhat between DTOA conditions (DTOA0: 132 ms; DTOA50: 138 ms; DTOA100: 122 ms; DTOA150: 141 ms) and were initially much higher in the DTOA0 condition than the remaining DTOA conditions (132-236 ms) (see Figure 6*D*). By ~250 ms of distractor processing, error rates were indistinguishable between DTOA conditions.

Taken together, these results show that although SRT modulated saccade metrics, excitatory visual onset responses were specifically related to distractor processing time, which were consistent with overall and OS estimates.

## Discussion

We non-invasively measured human saccade curvatures, endpoint deviations, saccade frequencies, and error proportions—indices of oculomotor excitation (Glimcher and Sparks, 1993; McPeek, 2006; McPeek et al., 2003; Munoz and Istvan, 1998; Port and Wurtz, 2003; Reingold and Stampe, 2002; Robinson, 1972; Robinson and Fuchs, 1969; White et al., 2012) and inhibition (Aizawa and Wurtz, 1998; Lee et al., 1988; McPeek and Keller, 2004; Munoz and Istvan, 1998; White et al., 2012)—as a function of time after the abrupt onset of a task-relevant, visually complex distractor. Critically, we ensured distractor feature processing and manipulated the visual similarity between targets and distractors. Based on previous physiological experiments, we hypothesized that distractor onsets would elicit an excitatory, feature-invariant visual onset response, followed by an inhibitory response encoding the object features.

On average, the latency of the visual onset response was ~135 ms for saccade curvature and ~110 ms for endpoint deviation. This 25 ms difference is consistent with saccade curvature as indicative of competing activity ~20-30 ms prior to saccade execution and endpoint deviation as indicative of competing activity at the time of saccade execution, demonstrating high reliability between these metrics and suggesting that the latency of visual onset responses was ~110 ms. Visual onset responses measured invasively in oculomotor substrates are typically much faster (reviewed by Boehnke and Munoz, 2008), but can increase as stimulus features require processing in higher areas of the ventral visual processing hierarchy (e.g., perceptual color space; White et al., 2009) or for extremely low luminance stimuli (e.g., ≤.05 cd/m^2^; Bell et al., 2006; Marino et al., 2015). As our stimuli were very high luminance (~125 cd/m^2^) and sufficiently complex to require processing in late stages of the ventral visual processing stream, the unusually long visual onset response latency of 110 ms we observed must reflect a delay in cortical feedforward projections into oculomotor substrates to accommodate sufficient cortical visual processing necessary for feature discrimination. As such, our results suggests that the contribution of the oculomotor substrates to feature-based target selection is limited to selecting the preprocessed cortical object representation with the highest activation as opposed to discriminating the target from distractor(s) on the basis of a local featural analysis. Strong corroboration of this account is the observation that error rates were not different from zero in the first ~150 ms after distractor onset and never occurred prior to 125 ms after distractor onset, suggesting that saccadic target selection was guided exclusively by the target representation during this epoch. A 40 ms delay between visual onset responses modulating saccade trajectories versus modulating saccade target choices suggests a buildup of the representational strength of the distractor until it was suprathreshold for eliciting saccades to the distractor, consistent with collicular visual onset bursts eliciting express saccades after sufficient buildup (Marino et al., 2015). We also observed a transient drop in the likelihood of making a saccade ~130 ms after distractor onset. This transient drop in saccade frequency occurs ~50-60 ms after a luminance flash on a saccade-to-target task (Reingold and Stampe, 2002), which further corroborates our account that the delayed visual onset response latency we observed relative to other experiments utilizing simpler visual features and task-demands, reflects the additional time required for sufficient cortical processing of stimuli to satisfy the current task demands.

We observed that the visual onset response, conventionally conceptualized as being strictly bottom-up and feature invariant (Boehnke and Munoz, 2008), was modulated by the behavioral relevance of the distractor: first, saccade endpoints were more biased towards the high similarity distractor than the intermediate similarity distractor 20 ms after the manifestation of the visual onset response (130 ms). Second, we observed that the excitatory epoch characterizing the visual onset response extended longer in time for the high similarity distractor than all other distractors as indicated by saccade curvature, endpoint deviation, and saccade frequency. Uniquely for the high similarity distractor, we observed two sequential excitatory epochs separated by ~80 ms, consistent with the physiological observation of a secondary visual onset burst in SCi VM cells (McPeek and Keller, 2002). Critically, the secondary excitatory epoch occurred prior to the inhibitory processing for the low and intermediate similarity distractors, which suggests that the protracted excitatory epoch encoding high similarity distractors does not merely reflect a delayed onset of inhibition. Since distractor identity could be decoded from the magnitude and duration of excitatory processing related to the distractor onset, this suggests that the oculomotor system dynamically receives preprocessed object representations from relevant cortical modules and encodes these objects as dynamically reweighted oculomotor vectors, as we have argued previously (Kehoe and Fallah, 2017; Kehoe et al., 2018a, 2018b).

Saccade curvature indicated that the magnitude of inhibition increased as distractors became increasingly dissimilar to the target, which we have observed previously using the same stimulus set (Kehoe et al., 2018a). This is the opposite pattern of results observed in previous behavioral studies of saccade curvature using opponent color singletons (Ludwig and Gilchrist, 2003; Mulckhuyse et al., 2009). This different pattern of results is likely related to methodological differences: as we utilized a perceptual discrimination here and previously (Kehoe et al., 2018a), other behavioral studies of the effects of visual similarity on saccade curvature utilized spatially guided saccades with task-irrelevant distractors (Ludwig and Gilchrist, 2003; Mulckhuyse et al., 2009). Perhaps on discrimination tasks, saccadic inhibition guides the target selection process itself and is therefore proportional to *perceptual confidence* (see Gold and Shadlen, 2000); while on spatially guided saccade tasks, saccadic inhibition guides saccadic accuracy to spatially preselected targets and is proportional to perceptual interference.

Erroneous saccades to the high similarity distractor persisted across time, while erroneous saccades to the intermediate and low similarity distractors gradually decreased to zero, suggesting that high similarity distractor representations were never successfully pruned from the decision process. As measured by saccade curvature, the onset of inhibition occurred first for the intermediate similarity distractor (215 ms), then the low similarity distractor (247 ms), and finally for the high similarity distractor (303 ms). Taken together, these observations are consistent with attentional facilitation of high similarity distractors within the featural focus of the attended target and intermediate distractors in the inhibitory annulus around the featural focus of the attended target, as would be predicted by the selective tuning model of attention (Tsotsos et al., 1995; Tsotsos, 2011) and a multidimensional (i.e., multi-featural) object-space (Kehoe et al., 2018a).

Splitting the data between the early, middle, and late blocks of the experiment revealed that the initial excitatory epoch, as measured by saccade curvature, was not temporally stable and immutable, but rather that it was reduced in the middle experimental blocks and was eventually extinguished in the late experimental blocks. In the middle blocks, there was an inhibitory response that immediately preceded the excitatory response. This observation suggests a strong influence of perceptual learning and executive function on saccade target selection processing, as executive processing learned the consistent latencies of excitatory projections into the oculomotor system to anticipatorily minimize distractor bias on target selection. Perhaps this anticipatory response was eventually well calibrated enough that the excitatory and anticipatory responses nullified one-another thus there was eventually no longer any evidence of either. On very similar perceptual discrimination saccade tasks, neuronal activity in executive area dorsal lateral prefrontal cortex was selective for both task type and object identity (Johnston and Everling, 2006) and relays this activity to cells in SCi (Johnston and Everling, 2006), which offers a plausible mechanism for the anticipatory response we observed. There was no evidence of visual onset response attenuation with the remaining saccade metrics. In fact, endpoint deviation, saccade frequency, and error proportion visual onset responses had approximately the same temporal properties as with the overall data across experimental blocks. We discuss this discrepancy in more detail below.

### Computational Modeling of Oculomotor Excitation and Inhibition

The current experiment demonstrates the versatility of this paradigm for non-invasively estimating the onset latency of oculomotor excitation and inhibition. There are several main differences between the current experiment and the original implementation (Kehoe and Fallah, 2017). First, using far more complicated, task-relevant stimuli increased the latency of the visual onset response by ~55 ms, consistent with visual onset burst latency differences between V1 and inferotemporal cortex (IT) (Nowak and Bullier, 1997) or V1 and V4 (Schmolesky, 1998). Similarly, we observed that inhibitory processing onset far later and accumulated far slower compared to inhibitory processing of task-irrelevant luminance- or color-modulated Gabor patches (Kehoe and Fallah, 2017). Second, we have utilized three additional behavioral metrics that are independent of saccade curvature, but that are all theoretically indicative of the same underlying oculomotor processes. These metrics provided consistent temporal estimates and have thus strengthened the validity of saccade curvature modeling to infer oculomotor excitation latencies. Third, to ensure that stimulus onset order did not provide useful target information, distractors onset prior to the target on half of the trials. However, SRTs were not different between DTOA conditions and there were strong effects of DTOA on target selection accuracy, suggesting that subjects prioritized speed over accuracy and were seemingly committed to their target choice. Furthermore, we analyzed directly whether visual onset responses were related to SRT or distractor processing time. Despite the fact that SRT modulated the visual onset response magnitude, we observed reliable visual onset responses with consistent latencies across DTOA conditions demonstrating that visual onset responses latencies are related to distractor processing time independently of SRT. As such, the current analyses can be interpreted as in the original paradigm: the competitive influence of a distractor over time while a saccade is concurrently being planned independently to a target.

### Putative Neural Mechanisms

Given the physiological spatiotemporal oculomotor interactions that elicit saccade curvature (Aizawa and Wurtz, 1998; McPeek, 2006; McPeek et al., 2003; Port and Wurtz, 2003; White et al., 2012) and shifted saccade endpoints (Fuchs and Robinson, 1969; Glimcher and Sparks, 1993; Lee et al., 1988; Robinson, 1972), distractor-related saccade curvature on our task must reflect the distractor-related neural activity level of VM cells ~20-30 ms prior to the saccade, while endpoint deviation must reflect neural activity levels of VM cells at the time of saccade initiation. If we therefore subtract 25 ms from our saccade curvature visual onset response latency estimate, this should correspond to the latency of visual onset bursts in the oculomotor substrates (SCi: McPeek et al., 2003; Port and Wurtz, 2003; White et al., 2012; and FEF: McPeek, 2006). On average, we observed a transient increase in positive saccade curvature with a latency of 135 ms. Therefore, the visual onset burst latencies of collicular VM cells eliciting this effect are 135 ms – 25 ms = 110 ms, exactly consistent with endpoint deviation. It seems unlikely that visual feedforward projections with this latency would originate from early vision, as parvocellular signals arrive in LGN 56 ms after stimulus onset and in layer 4Cβ of V1 by 67 ms (Schmolesky et al., 1998). Visual onset latencies in IT cells during passive viewing are between 100-110 ms (Nowak and Bullier, 1997), or as early as 80 ms during template-matching tasks where these signals differentiate targets from distractors 10 ms after onset (Miller et al., 1991). Given that IT is specialized for complex object discrimination (Miller et al., 1991), that there are direct projections from IT to the superior colliculus (Webster et al., 1993), and that visual onset latencies in IT during a similar template-matching discrimination task immediately precede the estimated time of the distractor-related visual onset response in SCi, we suspect that IT is the origin of the oculomotor object representations observed on the current task.

All four saccade metrics were indicative of excitatory oculomotor processing. However, three, presumably top-down, executively mediated inhibitory effects were only observed for saccade curvature: first, after the visual onset response, saccades increasingly began to deviate away from the distractor. The magnitude of this inhibitory response was modulated by target processing time (i.e., it was maximum for synchronous target-distractor onsets and was reduced as target lead time increased). Second, at the shortest possible SRTs (100-150 ms), saccade curvatures were consistently curved towards the distractor. Third, the saccade curvature visual onset response was gradually extinguished over the time course of the experimental session.

The distinguishing feature of saccade curvature, transience, provides some insight into unique saccade curvature effects. Oculomotor selection circuits are not strictly winner-take-all, as co-active saccade vectors elicit vector-weighted average movements (Fuchs and Robinson, 1969; Robinson, 1972). If co-activation is resolved before saccade initiation, then only an initial portion of the resultant saccade is vector-averaged but the saccade otherwise remains accurate, and the saccade trajectory is therefore curved (McPeek et al., 2003; McPeek, 2006; Port and Wurtz, 2003). Conversely, if co-activation persists to the time of saccade initiation, the entire saccade is vector-averaged (Glimcher and Sparks, 1993). As such, the emergence of unique saccade curvature effects suggests the emergence of transient modulation of the distractor and/or target vector-representations immediately preceding saccade execution, while attenuation of saccade curvature effects not apparent with other saccade metrics suggests the cessation of such transient modulation. Critically, the fact that this modulation is constrained to the narrow interval immediately preceding the saccade (White et al., 2012) highlights the top-down nature of such effects, as this temporally coincides with the saccadic go-signal, itself presumably related to the perceptual decision threshold (Ding and Gold, 2010). The shift from saccade curvature directed towards distractors and subsequently away from distractors observed as a function of SRT (Mulckhuyse et al., 2009; White et al., 2012) or distractor processing time (Kehoe and Fallah, 2017) and the attenuation of saccade curvature visual onset responses must then reflect a systematic shift in *how* saccades are triggered by the voluntary saccadic control system over the time course of individual trials and experimental sessions. For example, saccades may be triggered by direct excitatory input from dorsal-attentional cortices or indirectly from executive cortices via basal ganglian disinhibition (Hikosaka et al., 2000). Similarly, SNr contains both inhibitory and disinhibitory burst cells imposing concurrent suppressive and facilitatory effects on SCi (Shin and Sommer, 2010). The exact physiological mechanism for top-down effects apparent with saccade curvature can only be speculated given the emerging understanding of the voluntary saccade control system (see Basso and Sommer, 2012). However, it follows from the spatiotemporal interactions that elicit saccade curvature that systematic differences between interactive saccade trigger mechanisms must elicit unique saccade curvature effects. As such, these effects may be instantiated in a narrow window of time physiologically (McPeek, 2006; McPeek et al., 2003; Port and Wurtz, 2003; White et al., 2012), but apparent across an extended window of time behaviorally.

### Conclusion

We expanded on our paradigm in which human saccade metrics are non-invasively modeled as a function of time after abrupt distractor onset while a saccade is being independently planned to a target (Kehoe and Fallah, 2017) by utilizing visually complex, novel, and task-relevant stimuli that needed to be perceptually discriminated for successful target selection. We strengthened our neural interpretation of the data by using three additional behavioral metrics indicative of oculomotor excitation independent of saccade curvature and which gave consistent temporal estimates of visual onset response latencies. Our data show that the latencies of oculomotor visual onset responses elicited by complex objects are inconsistent with projections from early vision, suggesting instead that critical oculomotor substrates receive direct visual inputs from context-dependent cortical visual areas specialized for processing task-relevant features. Critically, we provide evidence that initial oculomotor excitatory responses can encode stimulus identity, contrary to influential views of oculomotor processing (Fecteau and Munoz, 2006). Stimuli were encoded according to perceptual confidence and we observed a strong role of executive perceptual learning in mediating these representations.

## Acknowledgements

We would like to thank our colleagues, past and present, in the Visual Perception and Attention lab for their thoughtful discussions and insights.

## References

Aizawa H, Wurtz RH. Reversible inactivation of monkey superior colliculus. I. Curvature of saccadic trajectory. J Neurophysiol 79: 2082–2096, 1998.

Basso MA, Sommer MA. Exploring the role of the substantia nigra pars reticulata in eye movements. Neuroscience 198, 205–212, 2012.

Bell AH, Meredith MA, Van Opstal AJ, Munoz DP. Stimulus intensity modifies saccadic reaction time and visual response latency in the superior colliculus. Exp Brain Res 174: 53–59, 2006.

Bichot NP, Schall JD. Effects of similarity and history on neural mechanisms of visual selection. Nat Neurosci 2: 549–554, 1999.

Boehnke SE, Munoz DP. On the importance of the transient visual response in the superior colliculus. Curr Opin Neurobiol 18: 544–551, 2008.

Carello CD, Krauzlis RJ. Manipulating intent: Evidence for a causal role of the superior colliculus in target selection. Neuron 43: 575–583, 2004.

Cavanaugh J, Wurtz RH. Subcortical modulation of attention counters change blindness. J Neurosci 24: 11236–11243, 2004.

Ding L, Gold JI. Caudate encodes multiple computations for perceptual decisions. J Neurosci 30, 15747–15759, 2010.

Dorris MC, Olivier E, Munoz DP. Competitive integration of visual and preparatory signals in the superior colliculus during saccadic programming. J Neurosci 27: 5053–5062, 2007.

Fecteau JH, Munoz DP. Salience, relevance, and firing: A priority map for target selection. Trends Cogn Sci 10: 382–390, 2006.

Fuchs AF, Robinson DA. Eye movements evoked by stimulation of frontal eye fields. J Neurophysiol 32: 637–648, 1969.

Glimcher PW, Sparks DL. Effects of low-frequency stimulation of the superior colliculus on spontaneous and visually guided saccades. J Neurophysiol 69: 953–964, 1993.

Gold JI, Shadlen MN. Representation of a perceptual decision in developing oculomotor commands. Nature 404: 390–394, 2000.

Hikosaka O, Takikawa Y, Kawagoe R. Role of the basal ganglia in the control of purposive saccadic eye movements. Physiol Rev 80, 953–978, 2000.

Horwitz GD, Newsome WT. Target selection for saccadic eye movements: Direction-selective visual responses in the superior colliculus. J Neurophysiol 86: 2527–2542, 2001.

Johnston K, Everling S. Neural activity in monkey prefrontal cortex is modulated by task context and behavioral instruction during delayed-match-to-sample and conditional prosaccade-antisaccade tasks. J Cogn Neurosci 18: 749–765, 2006.

Johnston K, Everling S. Task-relevant output signals are sent from monkey dorsolateral prefrontal cortex to the superior colliculus during a visuospatial working memory task. J Cogn Neurosci 21: 1023–1038, 2009.

Kehoe DH, Aybulut S, Fallah M. Higher-order, multifeatural object encoding by the oculomotor system. J Neurophysiol 120: 3042–3062, 2018a.

Kehoe DH, Fallah M. Rapid accumulation of inhibition accounts for saccades curved away from distractors. J Neurophysiol 118: 832–844, 2017.

Kehoe DH, Rahimi M, Fallah M. Perceptual color space representations in the oculomotor system are modulated by surround suppression and biased selection. Front Sys Neurosci 12:1, 2018b.

Lee C, Rohrer WH, Sparks DL. Population coding of saccadic eye movements by neurons in the superior colliculus. Nature 332: 357–360, 1988.

Lovejoy LP, Krauzlis RJ. Inactivation of superior colliculus impairs covert selection of signals for perceptual judgements. Nat Neurosci 13: 261–266, 2010.

Ludwig CJH, Gilchrist ID. Target similarity affects saccade curvature away from irrelevant onsets. Exp Brain Res 152: 60–69, 2003.

Marino RA, Levy R, Munoz DP. Linking express saccade occurance to stimulus properities and sensorimotor integration in the superior colliculus. J Neurophysiol 114: 879–892, 2015.

McPeek RM, Han JH, Keller EL. Competition between saccade goals in the superior colliculus produces saccade curvature. J Neurophysiol 89: 2577–2590, 2003.

McPeek RM, Keller EL. Deficits in saccade target selection after inactivation of superior colliculus. Nat Neurosci 7: 757–763, 2004.

McPeek RM, Keller EL. Saccade target selection in the superior colliculus during a visual search task. J Neurophysiol 88: 2019–2034, 2002.

McPeek RM. Incomplete suppression of distractor-related activity in the frontal eye field results in curved saccades. J Neurophysiol 96: 2699–2711, 2006.

Miller EK, Li L, Desimone R. Activity of neurons in anterior inferior temporal cortex during a short-term memory task. J Neurosci 13: 1460–1478, 1991.

Monosov IE, Sheinberg DL, Thompson KG. The effects of prefrontal cortex inactivation on object responses of single neurons in the inferotemporal cortex during visual search. J Neurosci 31: 15956–15961, 2011.

Moore T, Armstrong KM. Selective gating of visual signals by microstimulation of frontal cortex. Nature 421: 370–373, 2003.

Moore T, Fallah M. Control of eye movements and spatial attention. Pro Natl Acad Sci USA 98: 1273–1276, 2001.

Moore T, Fallah M. Microstimulation of the frontal eye field and its effects on covert spatial attention. J Neurophysiol 91: 152–162, 2004.

Mulckhuyse M, Van der Stigchel S, Theeuwes J. Early and late modulation of saccade deviation by target distractor similarity. J Neurophysiol 102: 1451–1458, 2009.

Müller JR, Philiastides MG, Newsome WT. Microstimulation of the superior colliculus focuses attention without moving the eyes. Pro Natl Acad Sci USA 102: 524–529, 2005.

Munoz DP, Istvan PJ. Lateral inhibitory interactions in the intermediate layers of the monkey superior colliculus. J Neurophysiol 79: 1193–1209, 1998.

Nowak LG, Bullier J. The timing of information transfer in the visual system. In: Cerebral Cortex: Extrastriate Cortex in Primates, edited by Rockland KS, Kaas JH, and Peters A. New York: Plenum Press, 1997, p. 205–241.

Palmer S. Structural aspects of visual similarity. Mem Cognit 6: 91–97, 1978.

Port NL, Wurtz RH. Sequential activity of simultaneously recorded neurons in the superior colliculus during curved saccades. J Neurophysiol 90: 1887–1903, 2003.

Reingold EM, Stampe DM. Saccadic inhibition in voluntary and reflexive saccades. J Cogn Neurosci 14: 371–388, 2002.

Robinson DA. Eye movements evoked by collicular stimulation in the alert monkey. Vision Res 12: 1795–1808, 1972.

Schall JD, Morel A, King DJ, Bullier J. Topography of visual cortex connections with frontal eye field in macaque: Convergence and segregation of processing streams. J Neurosci 15: 4464–4487, 1995.

Schmolesky MT, Wang Y, Hanes DP, Thompson KG, Leutgeb S, Schall JD, Veventhal G. Signal timing across the macaque visual system. J Neurophysiol 79: 3272–3278, 1998.

Shen K, Paré M. Neural basis of feature-based contextual effects on visual search behavior. Front Behav Neurosci 5: 1–12, 2012.

Shin S, Sommer MA. Activity of neurons in monkey globus pallidus during oculomotor behavior compared with that in substantia nigra pars reticulata. J Neurophysiol 103, 1874–1887, 2010.

Thompson KG, Bichot NP, Sato TR. Frontal eye field activity before visual search errors reveals the integration of bottom-up and top-down salience. J Neurophysiol 93: 337–351, 2005.

Tsotsos JK, Culhane SM, Wai WYK, Lai Y, Davis N, Nuflo F. Modeling visual attention via selective tuning. Artif Intell 78: 507–545, 1995.

Tsotsos JK. A Computational Perspective on Visual Attention. Cambridge MA: MIT Press, 2011.

Webster MJ, Bachevalier J, Ungerleider LG. Subcortical connections of inferior temporal areas TE and TEO in macaque monkeys. J Comp Neurol 335: 73–91, 1993.

White BJ, Boehnke SE, Marino RA, Itti L, Munoz DP. Color-related signals in the primate superior colliculus. J Neurosci 29: 12159–12166, 2009.

White BJ, Munoz DP. Superior colliculus. In: Oxford Handbook of Eye Movements, edited by Liversedge SP, Gilchrist I, and Everling S. New York: Oxford University Press, 2011, p. 195–213.

White BJ, Theeuwes J, Munoz DP. Interaction between visual- and goal-related neuronal signals on the trajectories of saccadic eye movements. J Cogn Neurosci 24: 707–717, 2012.

